# Dynamic Schlafen 11 (*SLFN11*) expression revealed by single-cell RNA sequencing of clonal human cancer cells

**DOI:** 10.64898/2026.10.01.755884

**Authors:** Daiki Taniyama, Yoshitaka Inoue, Fathi Elloumi, Yoo Sun Kim, Nitin Roper, Charlie Seibert, Michael Kelly, Alexei Lobanov, Mirit I Aladjem, Augustin Luna, Yves Pommier

**Affiliations:** Laboratory of Molecular Pharmacology and Developmental Therapeutics Branch, Center for Cancer Research, National Cancer Institute, National Institute of Health, Bethesda, MD 20892, USA; Department of Molecular Pathology, Graduate School of Biomedical and Health Sciences, Hiroshima University, Hiroshima, Japan; Computational Biology Branch, National Library of Medicine, National Institute of Health, Bethesda, MD 20892, USA; Department of Computer Science and Engineering, University of Minnesota, Minneapolis, MN 55455, USA; Single Cell Analysis Facility, Center for Cancer Research, National Cancer Institute, National Institute of Health, Bethesda, MD 20892, USA; Advanced Biomedical Computational Science, Frederick National Laboratory for Cancer Research, Frederick, MD, 21702, USA

## Abstract

Schlafen 11 (SLFN11) is a key determinant of sensitivity to DNA-damaging therapies in multiple cancers including leukemia, Ewing sarcoma, head and neck cancers, mesothelioma and small cell lung cancer (SCLC). Here, utilizing immunofluorescence microscopy and single-cell RNA sequencing we characterized SLFN11 expression across eight different human cancer cell lines and found cell-to-cell heterogenous expression both at the protein and RNA levels. Gene set enrichment and pathway analyses show that *SLFN11* is coexpressed with a distinct stress-response state and enhanced genome maintenance programs while being inversely correlated with protein translation pathways. Publicly available single-cell RNA-sequencing datasets of human SCLC tumors confirmed the heterogeneous distribution of *SLFN11* expression and confirmed its association with replication-related transcriptional programs. These findings indicate that heterogeneous SLFN11 expression reflects dynamic regulation at the single-cell level. They also underscore the limitations of bulk proteomic and transcriptome profiling and point to a potential reservoir of intrinsic chemotherapy resistance.

## Introduction

SLFN11 is one of the six members of the human innate-immune response Schlafen family (1–6). It also governs the sensitivity of cancer cells to a broad range of DNA-damaging chemotherapies targeting DNA replication (7–10). These include platinum derivatives (cisplatin and carboplatin), DNA polymerase inhibitors (gemcitabine and cytarabine), topoisomerase I (TOP1) inhibitors (camptothecin, irinotecan, topotecan, deruxtecan and exatecan), topoisomerase II (TOP2) inhibitors (etoposide and doxorubicin), Poly (ADPribose) polymerase (PARP) inhibitors (olaparib, niraparib, talazoparib and rucaparib) and antimetabolites (hydroxyurea) (6, 8, 9). Retrospective studies using patient tumor samples by immunohistochemistry show that elevated SLFN11 expression is correlated with favorable therapeutic responses to these DNA-damaging agents across cancer types (11–24).

SLFN11 expression is presently known to be driven under the control of the ETS transcription factors (25), the JAK-STAT (26) and interferon pathways (27) and to be suppressed epigenetically in approximately one third of cancer cell lines by promoter methylation (28), histone deacetylation (29) and by the polycomb repressive complex (30). Endogenous SLFN11 expression differs markedly among treatment-naive cancers, reflecting a finely balanced interplay between epigenetic and transcriptional regulation. Its basal silencing is largely driven by epigenetic mechanisms, including DNA hypermethylation and EZH2-dependent chromatin modifications (28, 30). Conversely, in Ewing sarcoma and leukemia, *SLFN11* is robustly expressed through direct transcriptional activation by oncogenic factors such as EWS-FLI1 (26, 29) or via intrinsic interferon signaling mediated by STAT1 (8). Despite such knowledge of *SLFN11* transcriptional regulation, the mechanisms governing the spatial and subcellular localization of SLFN11 remain largely elusive.

Recent studies using single-cell RNA sequencing (scRNA-seq) and histological analyses in patient samples have revealed substantial intratumoral heterogeneity in SLFN11 expression (31, 32). To our knowledge, this feature has not, until now been considered despite its potential for selecting resistant cellular clones following chemotherapy. scRNA-seq enables detailed characterization of the cellular components of the tumor microenvironment and tumor heterogeneity (31, 32). This technology is usually applied to dissect the complex multicellular architecture of diverse human cancers. By resolving transcriptomic profiles at single-cell resolution, these approaches have uncovered rare therapy-resistant subpopulations, mapped dynamic state transitions and elucidated intricate intercellular communication within tumors (33–35). Such detailed dissection of intratumoral heterogeneity is now essential for understanding disease progression and identifying novel therapeutic vulnerabilities. scRNA-seq analyses of SCLC have shown that SLFN11 is specifically enriched in POU2F3- and NEUROD1-driven tumor subpopulations, thereby supporting the use of precision therapies, including PARP inhibitors for these specific cell clusters (36).

Here, we first demonstrate that the expression of SLFN11 in cancer cell lines is heterogenous from cell to cell within clonal populations. This led us to utilize scRNA-seq to elucidate the genetic networks and pathways associated with the heterogenous expression of *SLFN11*. Pathway enrichment analyses revealed that *SLFN11* expression is coupled with DNA repair and replication stress responses, while being negatively associated with protein translation pathways. We further extended these observations to a publicly available scRNA-seq dataset of human SCLC, which revealed a continuous distribution of SLFN11 expression and a conserved association between SLFN11 expression and replication-related transcriptional programs in primary tumors. Together, these results define a transcriptomic framework coordinating SLFN11 expression with stress-response states.

## Materials and Methods

### Cell culture

U2OS, CCRF-CEM, DU145, human SCLC cell lines Colo668, DMS79, H1618, and H2107 were obtained from the American Type Culture Collection. All cell lines were maintained in RPMI-1640 (Corning #MT10040CM) supplemented with 10% fetal bovine serum and 1% penicillin–streptomycin. Doxycycline-inducible U2OS cells expressing SLFN11 were generated by lentiviral transduction with pLVX-TetOne-blasticidin-3X Flag-SLFN11. Forty-eight hours after infection, transduced cells were selected with blasticidin (10 µg/ml; Gibco). All experiments were performed using low-passage cells (<15 passages), and mycoplasma contamination was routinely monitored using the MycoAlert Mycoplasma Detection Kit (Lonza).

### Immunofluorescence staining and image analysis

Cells were fixed with 4% paraformaldehyde for 15 min and subsequently permeabilized with 0.5% Triton X-100/PBS for 15 min. After blocking with 5% BSA in PBS for 1 h, cells were incubated overnight at 4 °C with anti-SLFN11 antibody (Santa Cruz Biotechnology, D2) diluted 1:1000 in 5% BSA/PBS. After three washes with PBS, cells were incubated for 1 h with Alexa Fluor-conjugated anti-mouse secondary antibody (Invitrogen, A-11004) diluted 1:1000 in 5% BSA/PBST. Cells were then washed three times with PBS and mounted in VECTASHIELD Antifade Mounting Medium with DAPI (Vector Laboratories, H-1200). Fluorescence images were captured on a Nikon SoRa spinning-disk confocal microscope equipped with a 60× objective. Signal intensities in individual cells were quantified using ImageJ. The quantified values were used to generate single-cell signal intensity and co-localization plots, and the resulting data were visualized in GraphPad Prism 10 (GraphPad Software).

### Single-cell RNA sequencing, cell hashing, and data processing

Prior to loading, single-cell suspensions from each sample were multiplexed using TotalSeq-B anti-human Hashtag antibodies (HTO_1–HTO_8, BioLegend; clones LNH-94 and 2M2; cat. nos. 394631, 394633, 394635, 394637, 394639, 394641, 394643, and 394645). Specifically, cells were incubated with the respective hashtag antibodies for 30 min on ice, followed by extensive washing to remove unbound antibodies. Pooled cell samples were then assessed for concentration and viability using a Luna-FX7 fluorescent cell counter. Cells were loaded onto the 10x Genomics Chromium Connect automated platform at a concentration of 2,000 cells/µL, aiming for a target recovery of 20,000 cells per lane. Both scRNA-seq and hash-tag oligonucleotide (HTO) libraries were co-prepared using the Chromium Single Cell 3′Gene Expression v3.1 kit with Feature Barcode technology and TotalSeq-B reagents, in strict accordance with the manufacturer’s instructions. Libraries were sequenced on an Illumina NovaSeq 6000 S1 flow cell utilizing a custom read configuration (28 bp for read 1, 10 bp for index 1, 10 bp for index 2, and 90 bp for read 2). Following base calling with RTA v3.4.4, raw sequencing data were demultiplexed and aligned to the GRCh38-2020-A human reference genome using Cell Ranger v7.2.0 (STAR 2.7.2a). Concurrently, feature barcode reads were mapped to their respective TotalSeq-B hashtag sequences (HTO_1–HTO_8) to generate the single-cell count matrices utilized for all subsequent downstream analyses.

### Quality control

Gene-barcode matrices were processed using a Seurat-based workflow (v. 4.1.1) (37). HTO counts were normalized using centered log-ratio (CLR) normalization and demultiplexed with MULTIseqDemux using automatic thresholding (autoThresh = TRUE). Cells classified as negative or cross-sample doublets were removed. Quality-control (QC) filtering was then performed to exclude low-quality cells, empty droplets, and transcriptomic doublets. Genes detected in fewer than 20 cells were removed. Cells were retained only if the number of detected genes (nFeature_RNA) and UMIs (nCount_RNA) were within five median absolute deviations (MADs) of the sample median. Cells with >25% mitochondrial reads were removed. Cells with low library complexity, defined using a lower cutoff of five MADs below the sample median for log10GenesPerUMI, were also removed. In addition, cells with outlier percentages of counts contributed by the top 20 expressed genes were filtered using five-MAD cutoffs. Transcriptomic doublets were identified with scDblFinder, and only cells classified as singlets were retained for downstream normalization, dimensionality reduction, and clustering. Filtered cells were normalized with SCTransform, followed by variable-feature selection, PCA, UMAP/t-SNE visualization, and graph-based clustering. After quality control, the final dataset contained 9,307 single cells across eight cell-line datasets. Each expression matrix contained 27,491 gene features. The scRNA-seq data generated in this study have been deposited in the Gene Expression Omnibus (GEO) database under accession code GSE336003.

### Gene filtering and expression analysis

For each cell line, genes were filtered based on their expression range across cells to prioritize those exhibiting substantial variability. Specifically, genes within the top 25% ranked by expression range (defined as the difference between the maximum and minimum expression across cells) were retained. If SLFN11 was not included in this subset, the filtering threshold was set to the expression range of SLFN11, and all genes with an expression range greater than or equal to this value were retained.

Pearson correlation coefficients between SLFN11 and the filtered genes were then calculated, and genes showing significant correlation (p < 0.05) were ranked based on their correlation coefficients.

### Imputation

To mitigate the impact of dropout events inherent to scRNA-seq data, we performed data imputation using the MAGIC (Markov Affinity-based Graph Imputation of Cells) algorithm (38). Dropout events arise primarily due to limited mRNA capture efficiency and stochastic detection of lowly expressed transcripts (39). MAGIC reconstructs gene expression values by leveraging similarities between cells in a diffusion space, thereby improving the continuity of gene expression signals across transcriptionally similar cells. Both the original and imputed expression matrices were used for downstream investigations to improve the robustness of correlation-based analyses, particularly for genes with low expression levels.

### Pathway analysis

To identify biological processes associated with SLFN11 expression across various cancer cell lines, we performed Gene Set Enrichment Analysis (GSEA) (40). We used the GSEApy library (v1.1.11) (41) in preranked mode with the Reactome pathway database (42) (MSigDB collection (43) c2.cp.reactome.v2025.1.Hs.symbols). Significantly enriched pathways were defined using a nominal P value (NOM P) < 0.05, as part of an exploratory analysis.

### Patient-derived single-cell RNA sequencing analysis

Publicly available scRNA-seq data from primary SCLC epithelial cells were obtained from the Human Tumor Atlas Network (HTAN) dataset “Single-cell profiling reveals novel tumor and myeloid subpopulations in small cell lung cancer” (44). The dataset comprised 54,313 cells from 19 patients. Patient-level *SLFN11* expression was quantified using normalized, log-transformed expression values, and the five patients with the highest mean *SLFN11* expression were selected for downstream analyses. Gene filtering, correlation analyses, and pathway enrichment analyses were subsequently performed as described above. The requirement for institutional review board (IRB) approval was waived because this study utilized publicly available, de-identified scRNA-seq datasets from the HTAN. Informed consent was obtained by the original investigators of the primary study, and all data were analyzed anonymously.

## Results

### Broad fluctuation of SLFN11 protein expression among clonal tumor cells

Immunofluorescence microscopy was performed to assess cell-to-cell variability in SLFN11 protein expression within six human cancer cell lines known to express *SLFN11* (8, 45, 46) (https://discover.nci.nih.gov/) (47): leukemia CCRF-CEM, prostate cancer DU145, and four small cell lung cancer (SCLC) cell lines (Colo668, DMS79, H1618, and H2107). SLFN11-negative control osteosarcoma cells (U2OS) were also included as well as its doxycycline *SLFN11*-inducible counterpart (Figure 1A-B). Both the *SLFN11*-inducible and the cells endogenously expressing *SLFN11* expression exhibited marked heterogeneity at the single-cell level.

**Fig 1.**
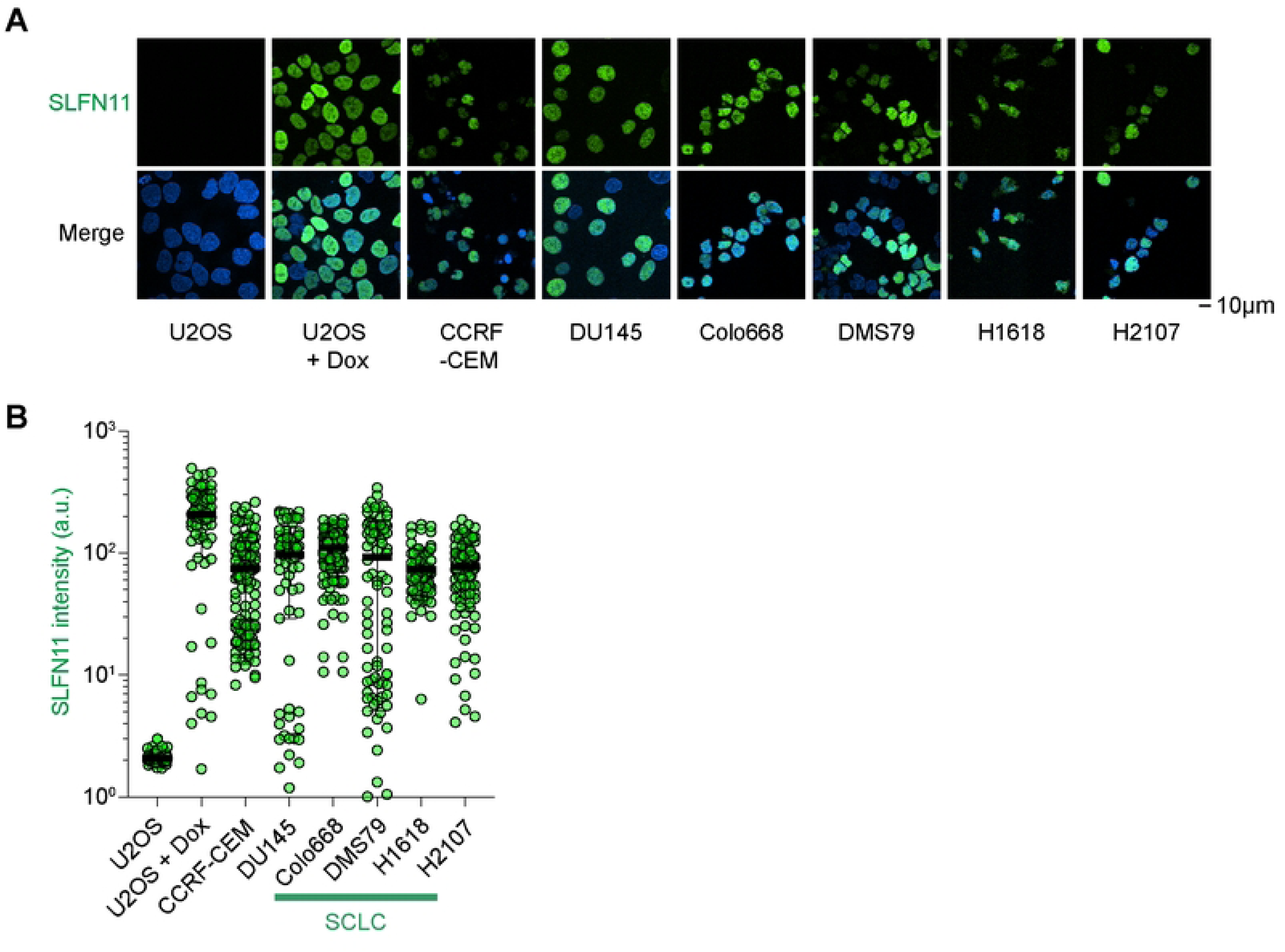
SLFN11 exhibits dynamic and bimodal protein expression in human single cells. (A) Representative immunofluorescence images of SLFN11 expression in osteosarcoma U2OS cells, doxycycline (DOX)-inducible SLFN11-expressing U2OS cells, leukemia CCRF-CEM cells, prostate cancer DU145 cells, and the SCLC cell lines Colo668, DMS79, H1618, and H2107 cells. Scale bar: 10 µm. (B) Quantification of SLFN11 signals in individual cells for the indicated cell lines. Mean ± SD are shown (n = 71 – 184 cells per condition).

### Cellular fluctuation of *SLFN11* mRNA expression within cancer cell lines

To elucidate the determinants of SLFN11 expression fluctuations, we performed scRNA-seq in the eight cancer cell lines using the 10x Genomics platform with hashing antibody libraries followed by read alignment (STAR) and gene expression quantification (Figure 2A).

**Fig 2.**
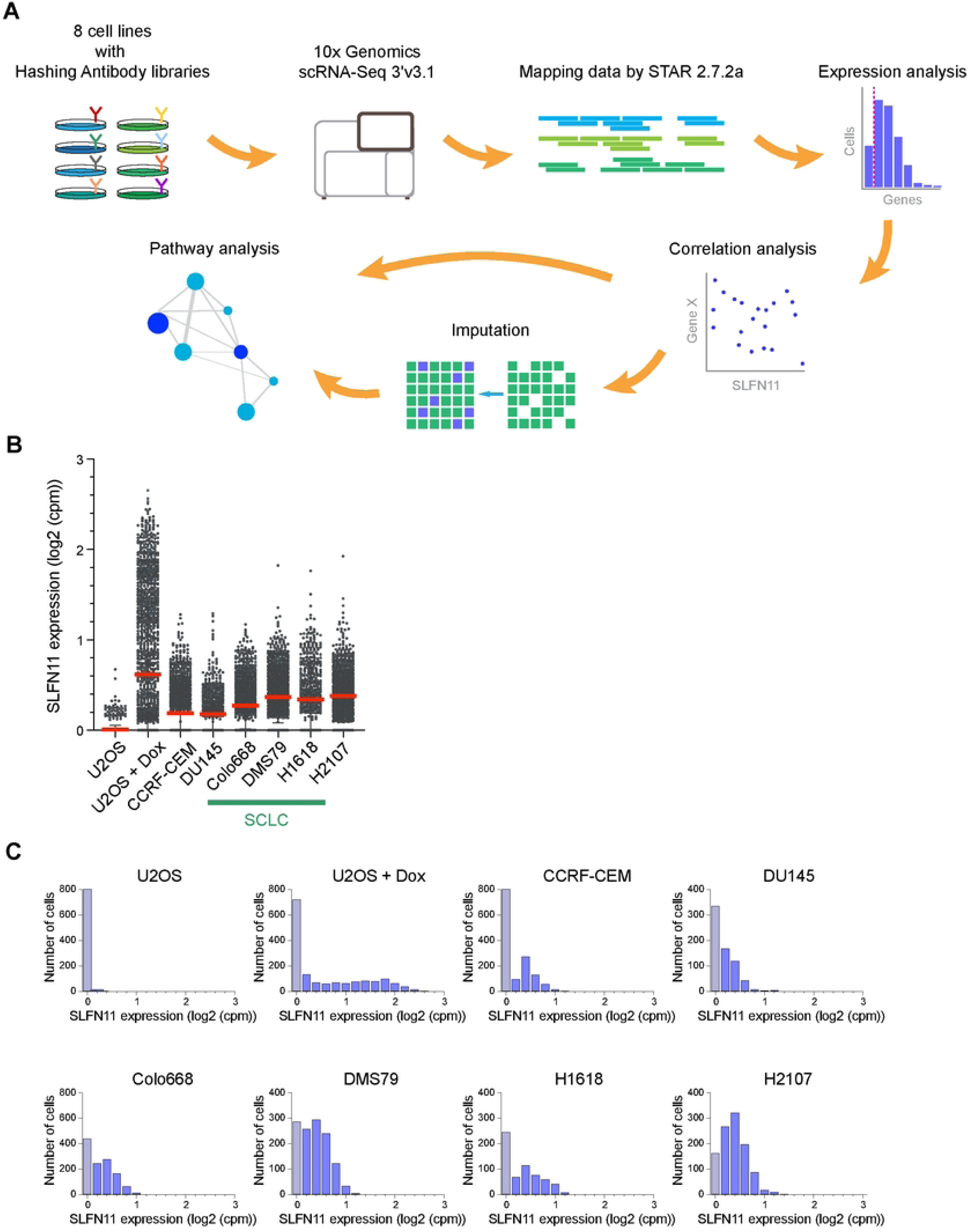
SLFN11 exhibits dynamic and continuous mRNA expression. (A) Schematic overview of the single cell RNA sequencing (scRNA-seq) and data analysis workflow. (B) scRNA-seq analysis of *SLFN11* mRNA expression across the eight cancer cell lines. Each dot represents *SLFN11* expression in an individual cell, with expression shown as log2 (counts per million, cpm). Mean expression levels ± SD are shown. (C) Distribution of *SLFN11* mRNA expression in individual cells for each cell line. The histograms show the frequency of cells across log 2 (cpm) expression bins, highlighting continuous variability at the mRNA level.

Consistent with our immunofluorescence results (see Figure 1), scRNA-seq showed heterogeneity in *SLFN11* expression across individual cells (Figure 2B-C). The *SLFN11*-negative U2OS cells, which served as negative control, showed minimal expression, whereas induction of *SLFN11* in U2OS + Dox resulted in a broad distribution of *SLFN11* expression levels. All *SLFN11*-expressing cell lines showed marked heterogeneity at the single-cell level. Hematopoietic (CCRF-CEM) and prostate (DU145) cancer cells exhibited predominantly low to intermediate average *SLFN11* expression with a subset of higher-expressing cells. The SCLC cell lines (Colo668, DMS79, H1618 and H2107) also displayed heterogeneous expression with generally high *SLFN11* expression. Notably, a fraction of cells consistently exhibited low or undetectable *SLFN11* expression even within *SLFN11*-high populations. These findings demonstrate extensive *SLFN11* cell-to-cell heterogeneity at the RNA level across diverse cancer cell lines from different histological types.

### Identification of genes co-expressed with *SLFN11*

To identify genes associated with *SLFN11* expression within individual cells, we performed correlation analyses between *SLFN11* and all expressed genes at the single-cell level (Figure 3). Genes were stratified by expression level, and those exhibiting a broad dynamic range (top 25%) were selected to capture heterogeneity paralleling *SLFN11* expression.

**Fig 3.**
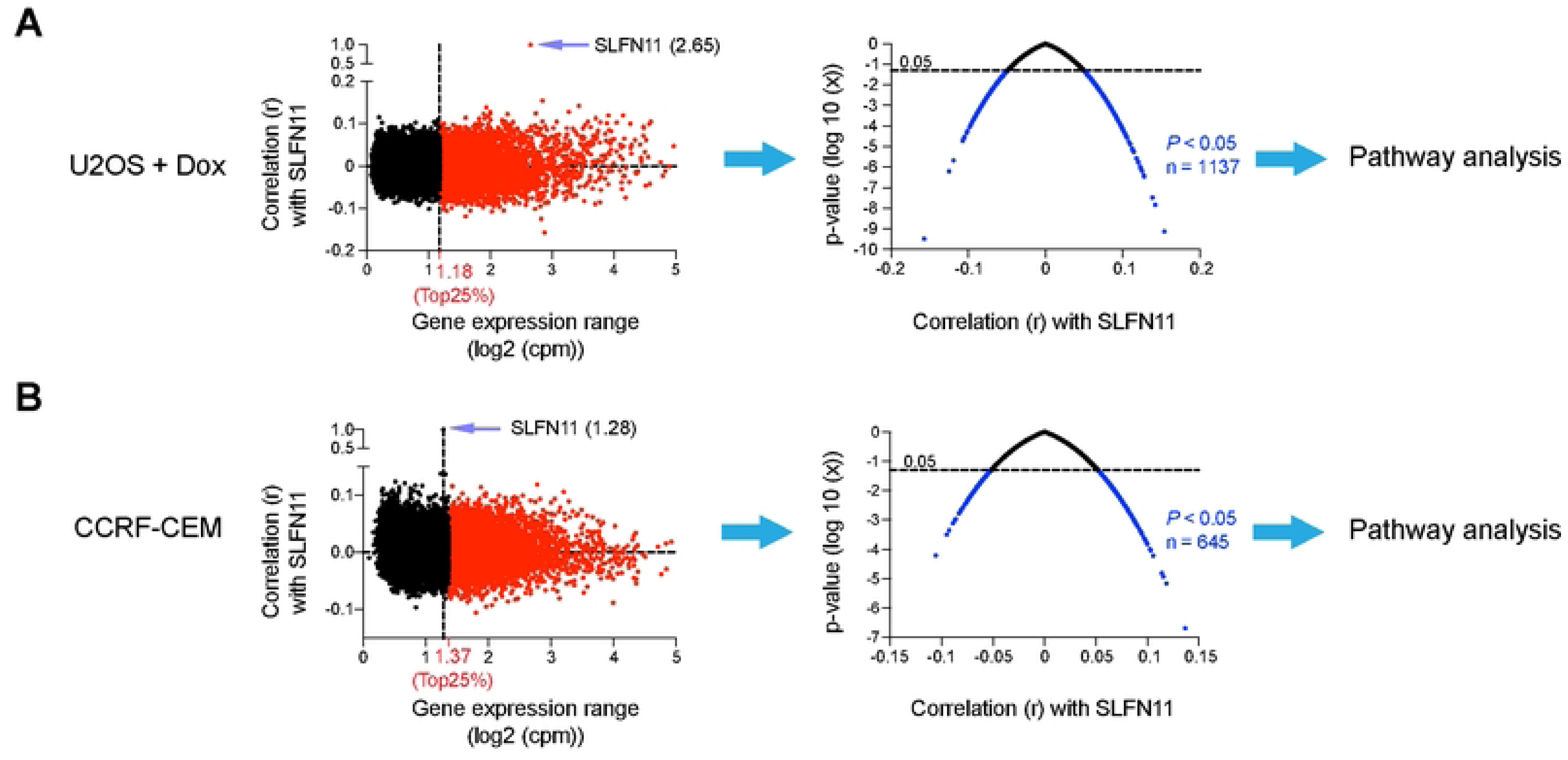
Identification of genes with a broad dynamic range of expression associated with SLFN11 expression. (A) Left: Genes expressed with a broad dynamic range (top 25%, shown in red) were selected. (B) as the *SLFN11* expression range (1.28) was below the top 25% threshold (1.37), the cutoff was set to the *SLFN11* range, and genes meeting or exceeding this value were analyzed. Right: statistical significance of correlations shown as *P* values (−log10 scale) plotted against correlation coefficients. Significantly associated genes (*P* < 0.05) are highlighted and subjected to pathway analysis. Data are shown for U2OS + Dox (A) and CCRF-CEM (B) cells.

In U2OS + Dox cells, *SLFN11* expression showed a wide expression range (2.65), placing it within the top 25% fluctuating genes. Statistical filtering (P < 0.05) identified 1,137 genes significantly associated with *SLFN11* expression (Figure 3A). In CCRF-CEM cells, *SLFN11* exhibited a narrower expression range (1.28), below the top 25% threshold (1.37); therefore, genes with expression ranges exceeding that of *SLFN11* were selected for correlation analysis. This approach identified 645 genes significantly associated with *SLFN11* expression (Figure 3B). In both models, the distribution of correlation coefficients was centered near zero but exhibited significant deviation at both positive and negative extremes. These *SLFN11*-associated gene sets were subsequently subjected to pathway analysis to uncover underlying biological processes.

Single-cell RNA-seq data are characterized by pervasive zero-inflation, which can obscure downstream analyses. To mitigate this effect, we applied imputation and performed correlation analyses before and after correction.

Across all cell lines, correlation analyses using raw data identified genes with modest associations with *SLFN11* expression, whereas imputation revealed significant and coherent expression patterns at the single-cell level, indicating improved recovery of underlying biological signals (Figures 4 and S1).

**Fig 4.**
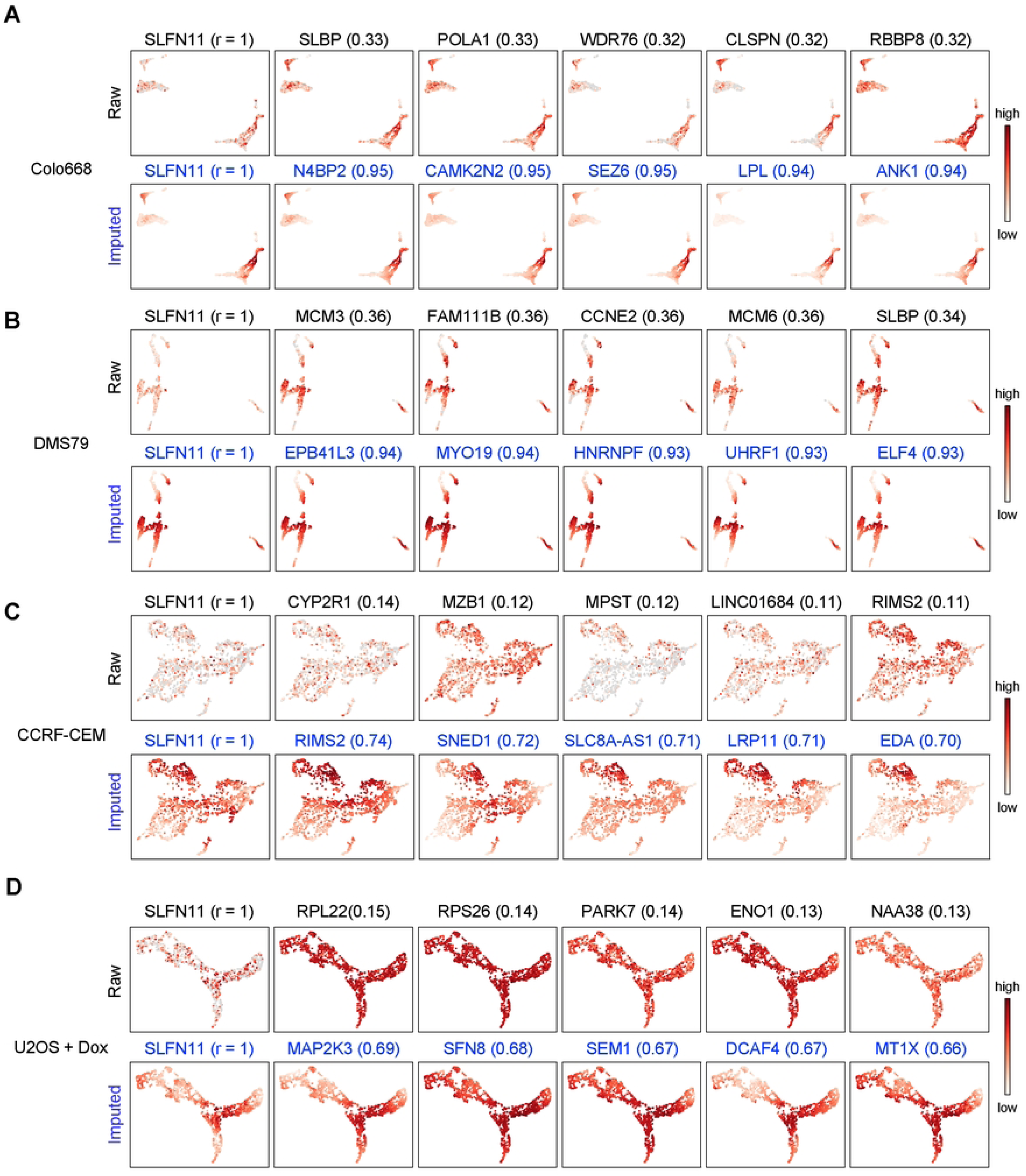
*SLFN11*-correlated gene networks. Two-dimensional UMAP projections of single cells from Colo668 (A), DMS79 (B), CCRF-CEM (C) and U2OS + Dox (D). *SLFN11* expression and the top five genes most correlated with *SLFN11* in each cell line are shown. For each panel, the top row shows raw expression and the bottom row imputed expression. *SLFN11* is shown to the left of each row as the reference gene (r = 1), followed by the highest-correlated genes identified from the corresponding raw or imputed data. Values in parentheses indicate the Pearson correlation coefficient (r) with *SLFN11* expression across single cells within each cell line. Color scale denotes relative expression from low (white) to high (red).

In the SCLC cell lines (Colo668, DMS79, H1618 and H2107), raw data revealed only modest correlations between *SLFN11* expression and a limited set of genes, including several involved in DNA replication and cell-cycle regulation (*SLBP*, *POLA1*, *CLSPN*, *MCM3* and *CCNE2*). Imputation strengthened these associations and uncovered more coherent, cell line-specific gene modules, extending beyond proliferation-related genes to include factors implicated in transcription, RNA processing and signaling *(N4BP2, CAMK2N2, SEZ6, LPL, ANK1, EPB41L3, MYO19, HNRNPF, UHRF1*, *ELF4*, *RIMS2*, *SNED1*, *SLC8A-AS1*, *LRP11* and *ED*A).

In non-SCLC cell lines (U2OS + Dox, CCRF-CEM and DU145), correlation analyses of raw data identified genes modestly associated with *SLFN11*, including regulators of neuroendocrine differentiation and metabolic adaptation (*RIMS2, SYNPR-AS1*, *MSX1, PARK7, MPST,* and *ENO1*). Imputation strengthened these associations and enhanced the coherence of expression patterns, reinforcing the link between *SLFN11* expression and neuroendocrine differentiation programs. Collectively, these findings demonstrate that imputation enhances the detection and interpretability of *SLFN11*-associated gene networks across diverse cancer cell types.

### Common enriched pathways ranked by normalized enrichment score (NES)

To define the biological pathways associated with *SLFN11* expression, we performed pathway enrichment analyses based on *SLFN11*-correlated genes across all cell lines with pathways ranked by normalized enrichment score (NES) (Figure 5). For each cell line, the top 10 positively and bottom 10 negatively enriched pathways (NOM P < 0.05) were extracted, and pathways were ranked by NES in each cell line. Those consistently highly ranked across cell lines were prioritized based on cumulative rank.

**Fig 5.**
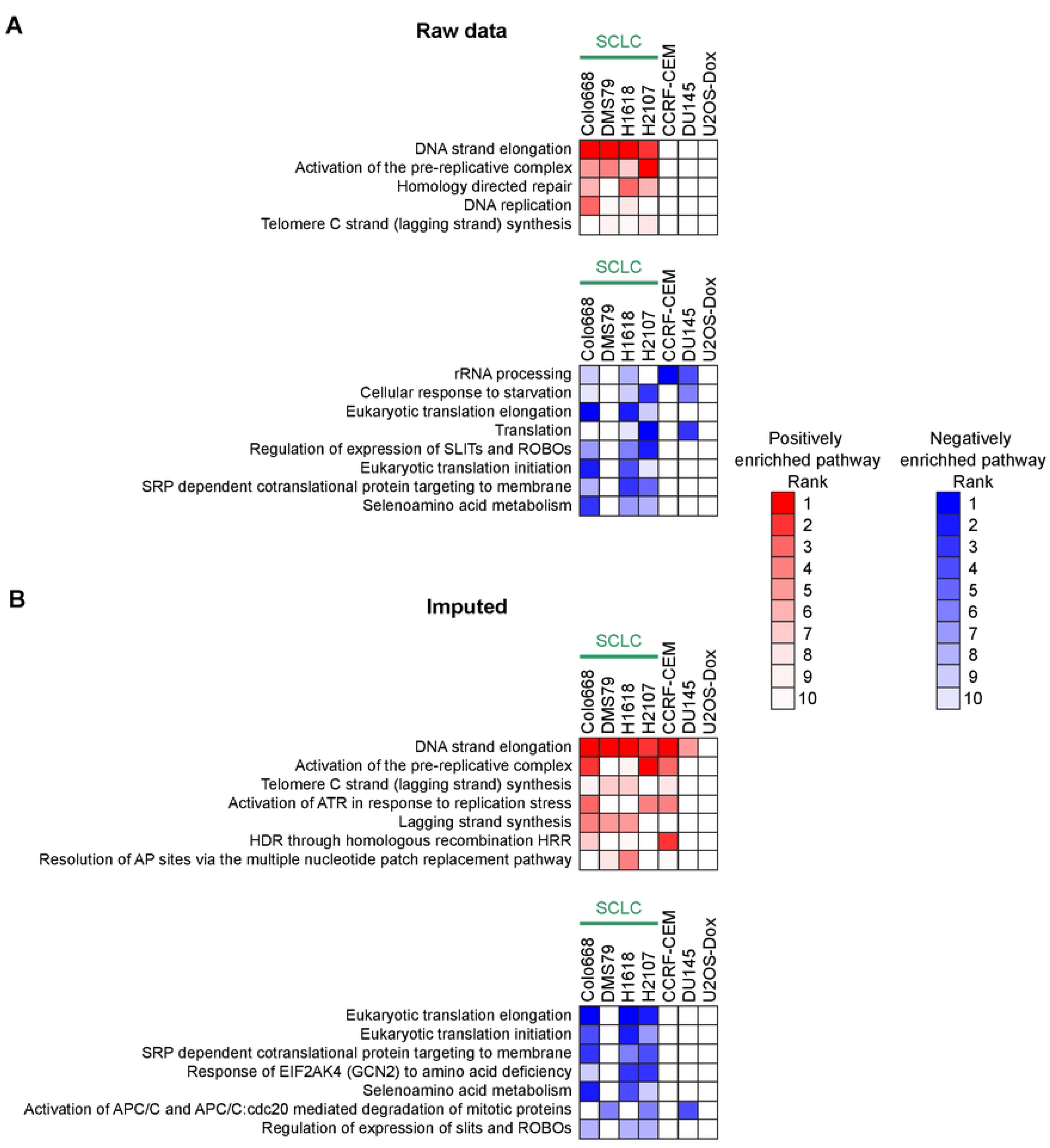
Pre-ranked GSEA reveals *SLFN11*-associated pathways across cell lines. The heat-maps show the pathways identified by pre-ranked GSEA in each cell line using genes ranked by their correlation with *SLFN11* expression in raw and imputed scRNA expression data (panels (A) & (B), respectively). Columns correspond to the indicated cell lines. Color intensity denotes pathway rank ordered by normalized enrichment score (NES) within each cell line (1, strongest enrichment; 10, weakest). White indicates that the pathway was not among the top 10-ranked pathways in that cell line. In raw data, positively-enriched pathways were mainly related to DNA replication and repair and were most consistently observed in the SCLC cell lines, whereas negatively enriched pathways were associated with protein translation and RNA processing and were more variable across cell lines. Imputation increased the consistency and specificity of these enrichment patterns.

Using the raw data, positively enriched pathways were predominantly related to DNA replication and repair, including DNA strand elongation, activation of the pre-replicative complex and homology-directed repair, with enrichment most consistently observed in SCLC cell lines. Negatively enriched pathways were largely associated with translation and RNA processing, including eukaryotic translation elongation and rRNA processing, although these signals were variable across cell lines.

Imputation enhanced both the consistency and specificity of pathway enrichment. Positively enriched pathways, ranked by NES, were reinforced and expanded to include replication stress response pathways, such as ATR activation and lagging-strand DNA synthesis, as well as homologous recombination repair. These pathways were consistently enriched across SCLC models but were also detected in the non-SCLC cell lines. Conversely, negatively enriched pathways identified after imputation showed stronger and more uniform enrichment when ranked by NES, prominently highlighting translation-related processes, including translation initiation, elongation and SRP (Signal Recognition Particle)-dependent protein targeting. Additional pathways related to cellular stress responses, such as EIF2AK4-mediated amino acid deficiency response, were also revealed following imputation.

Overall, imputation improved the resolution and reproducibility of pathway-level signals, reinforcing the association between *SLFN11* expression and DNA replication and repair programs, while uncovering coordinated suppression of translational machinery across diverse cancer cell types.

### High *SLFN11* expression is associated with DNA replication and replication stress response programs in human SCLC tumors

To extend our findings obtained in SCLC cell lines, we analyzed a publicly available scRNA-seq dataset of human SCLC tumors (44). Among 19 patient samples, the five samples with the highest mean *SLFN11* expression were selected for analysis. Consistent with our observations in the cell lines, *SLFN11* expression exhibited a continuous distribution across individual tumor cells rather than forming discrete *SLFN11*-positive and -negative populations (Figures 6A and S2).

**Fig 6.**
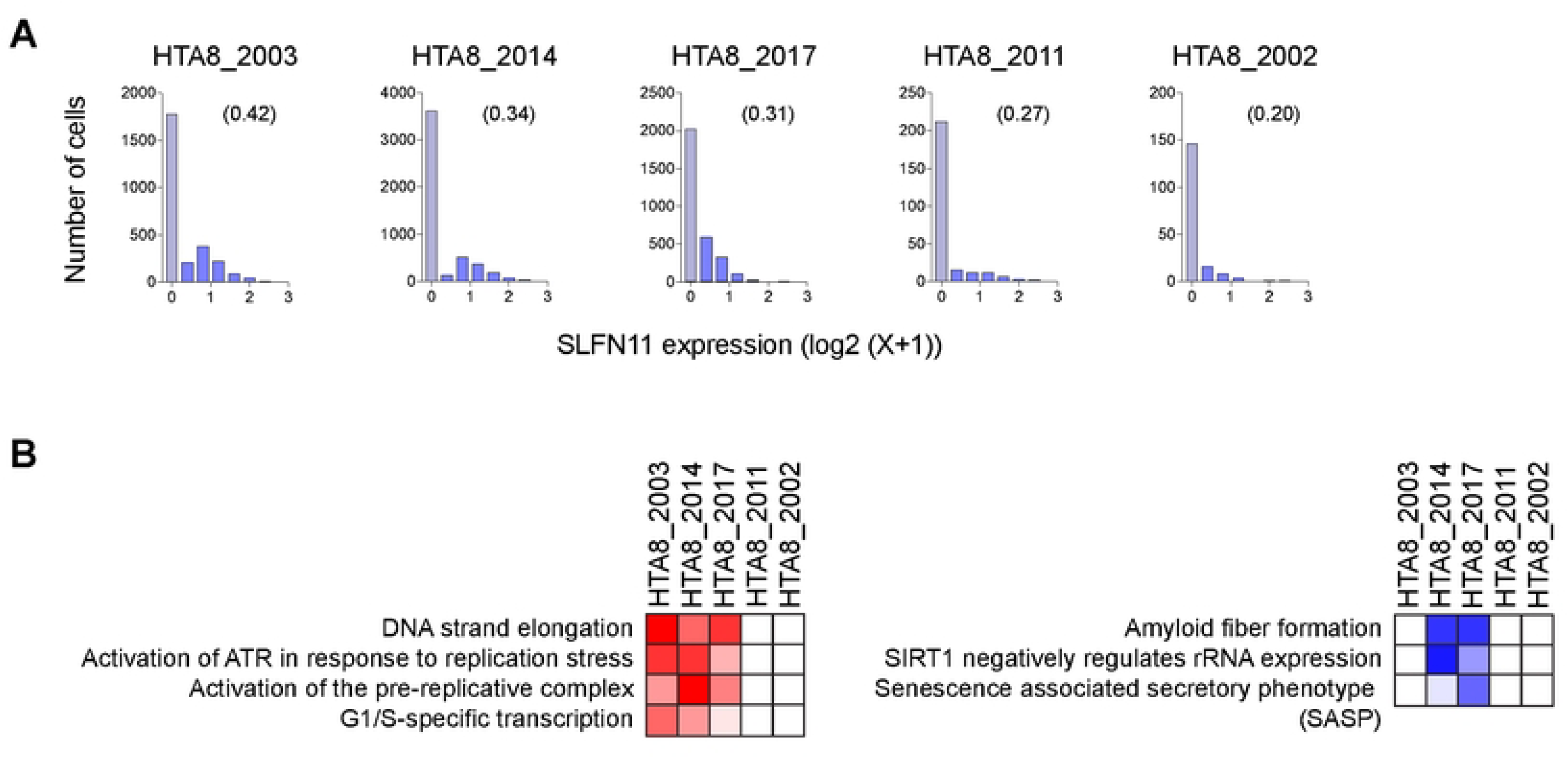
Heterogeneous SLFN11 expression and associated transcriptional programs in human SCLC samples. (A) Histograms showing the distribution of single-cell SLFN11 expression across individual SCLC samples. Publicly available scRNA-seq datasets were analyzed, and the five samples with the highest mean SLFN11 expression were selected from a cohort of 19 patients. *SLFN11* expression is shown as log2(X + 1), where X represents normalized transcript counts. Numbers in parentheses indicate the mean SLFN11 expression level for each sample. (B) Heat maps showing the pathways identified by pre-ranked GSEA in each sample using genes ranked by their correlation with SLFN11 expression in raw scRNA expression data. Columns correspond to the indicated samples. Color intensity reflects pathway ranking based on NES, as shown in Figure 5. Samples with high *SLFN11* expression were associated with enrichment of DNA replication–related pathways, including DNA strand elongation, activation of ATR in response to replication stress, activation of the pre-replicative complex, and G1/S-specific transcription. In contrast, pathways related to amyloid fiber formation, SIRT1― mediated repression of rRNA expression, and senescence associated secretory phenotype were negatively enriched.

Using the same filtering strategy applied to the cell line datasets, we performed pre-ranked GSEA based on genes correlated with *SLFN11* expression in each sample. Pathways positively associated with *SLFN11* expression included DNA strand elongation, activation of ATR in response to replication stress, activation of the pre-replicative complex, and G1/S-specific transcription (Figure 6B), closely recapitulating the transcriptional programs identified in SCLC cell lines. In contrast, negatively associated pathways included amyloid fiber formation, SIRT1-mediated repression of rRNA expression, and senescence associated secretory phenotype. These differed from the negatively enriched pathways identified in cell lines. Together, these findings support a conserved association between *SLFN11* expression and replication-related transcriptional programs in both SCLC cell lines and primary human tumors, while suggesting greater heterogeneity in pathways negatively correlated with *SLFN11* expression in patient samples.

## Discussion

This study demonstrates SLFN11 expression heterogeneity at the single-cell level and provides a framework for understanding how SLFN11 expression is integrated with transcriptional programs across diverse cancer types. Although bulk RNAseq and immunohistochemistry analyses of tumor samples show that SLFN11 is a well-established determinant of sensitivity to DNA-damaging therapies, here we demonstrate that its expression is highly dynamic at the single-cell level, including in cell lines commonly used to study SLFN11 molecular biology (7, 8, 48–51).

The pronounced cell-to-cell variability in SLFN11 expression observed in our scRNA-seq analyses highlights the intrinsically dynamic gene expression programs within individual cancer cells. This variability was observed under basal conditions in the absence of exogenous stress or drug treatment, indicating that SLFN11 expression is independent of acute replication stress induction. This is consistent with the fact that SLFN11 transcripts and protein levels are not induced by chemotherapies (7, 9). This feature bears conceptual resemblance to the well-characterized heterogeneity of p53; however, the underlying mechanisms are fundamentally distinct. p53 heterogeneity arises from pulsatile dynamics at the protein level, driven by an MDM2-mediated negative feedback loop that generates oscillatory responses following DNA damage (52, 53). In contrast, *SLFN11* expression exhibits a continuous distribution at the mRNA level across most cell lines, in contrast to the more binary patterns observed by immunofluorescence. These findings suggest that SLFN11 fluctuations might not be governed by feedback-driven oscillations but instead reflect differences in transcriptional activity and protein-level regulation across cells, supporting a model in which SLFN11 expression reflects dynamic cellular states that may contribute to variability in therapeutic response (30, 54).

Consistent with this view, *SLFN11* gene expression has been reported to be regulated epigenetically by DNA methylation (29, 55), histone acetylation (56) and chromatin remodeling (30), which could underlie the diverse transcriptional states observed at the single-cell level. The observation of bimodal protein abundance suggests that this variability may be further amplified or filtered by post-transcriptional and/or post-translational control, although the molecular basis of this regulation remains unresolved. Importantly, recent work has broadened the functional landscape of SLFN11 by implicating SLFN11 in proteotoxic stress regulation where loss of SLFN11 induces proteotoxic stress and renders cancer cells vulnerable to ubiquitin-activating enzyme inhibition (57). These observations raise the possibility that SLFN11 functions not only as a determinant of genome maintenance but also as a broader node in cellular stress adaptation, with heterogeneous expression potentially shaping multiple stress-responsive phenotypes.

Our analyses reveal that *SLFN11* expression is embedded within coordinated gene networks enriched for DNA replication and cell cycle across the seven human cancer cell lines studied here. These findings are in line with the established role of SLFN11 in replication stress responses and reinforce SLFN11’s functional coupling with proliferative cellular states (7, 8, 58). Similar gene networks were identified in both raw and imputed datasets, supporting the robustness of these associations. Pathway enrichment analyses show consistent enrichment of DNA replication, repair and replication stress response pathways, including ATR signaling and homologous recombination. In contrast, translation-related pathways were negatively associated with SLFN11 expression, suggesting that SLFN11 expression is linked to a distinct stress-response state characterized by enhanced genome maintenance and reduced translational activity.

In SCLC cell lines, we find that the *SLFN11-*associated gene modules extend beyond canonical replication pathways to include genes involved in transcription, RNA processing and signaling, indicating that *SLFN11*-associated programs extend beyond replication-related processes and may reflect broader cellular states such as protein synthesis (59–61). In non-SCLC cell lines, SLFN11-correlated genes were enriched for neuronal and neuroendocrine transcriptional programs. These findings suggest a previously underappreciated association between *SLFN11* expression and neuroendocrine cellular states.

To extend our cell-line observations, we interrogated a publicly available human SCLC single-cell atlas and found that the association between *SLFN11* expression and replication-related transcriptional programs was also evident in primary tumors. Notably, *SLFN11* expression was continuously distributed across patient samples rather than segregating into discrete *SLFN11*-positive and -negative populations. In the randomized phase II trial of temozolomide plus veliparib in relapsed SCLC, approximately 48% of tumors were classified as SLFN11-positive using an H-score cutoff of ≥1, and SLFN11 positivity was associated with improved clinical outcomes (18). Consistent with this prevalence, a mean *SLFN11* expression threshold of 0.1 identified approximately half of the tumors in the present single-cell cohort. However, among the five tumors with the highest *SLFN11* expression (mean expression >0.2), replication-associated transcriptional programs were consistently observed only in HTA8_2003, HTA8_2017, and HTA8_2014. These findings suggest that biologically meaningful heterogeneity exists among tumors that would otherwise be grouped together as SLFN11-positive.

Collectively, these observations support a model in which SLFN11 expression functions as a continuous rather than strictly dichotomous variable. Thus, binary classification based solely on SLFN11 positivity may not fully capture the biological diversity associated with SLFN11 expression. While the relationship between these transcriptional states and therapeutic response remains to be established, our data suggest that quantitative differences in SLFN11 expression may reflect distinct biological states associated with varying degrees of replication-associated activity.

Clinically, these results support the use of SLFN11 as a functional biomarker of replication stress–associated therapeutic vulnerability. However, the presence of heterogeneity within SLFN11-high populations suggests that bulk measurements may overestimate therapeutic responsiveness. The identification of SLFN11-low subpopulations further highlights the potential for intrinsic resistance and underscores the need for therapeutic strategies that can target or overcome these cells (11, 12, 30, 56). Together, these findings provide a refined framework for interpreting SLFN11 expression in the context of tumor heterogeneity and therapeutic response.

Our study has several limitations. First, the mechanistic and functional analyses were primarily performed in a limited panel of cancer cell lines and were enriched for SLFN11-high models. Although analysis of a publicly available SCLC single-cell dataset supported the association between SLFN11 expression and replication-related transcriptional programs in primary tumors, these observations were derived from retrospective transcriptomic analyses and were not linked to treatment outcomes. Second, while our data suggest that quantitative differences in SLFN11 expression may define biologically distinct tumor states, the relationship between these states and therapeutic responsiveness remains to be determined. Finally, although cell lines provide a controlled system for dissecting intrinsic heterogeneity, they do not recapitulate the full complexity of the tumor microenvironment. Therefore, further studies using patient-derived models and clinically annotated cohorts will be required to establish the biological and therapeutic significance of SLFN11-associated tumor heterogeneity.

## Acknowledgements

The contributions of the NIH author(s) are considered Works of the United States Government. The findings and conclusions presented in this paper are those of the author(s) and do not necessarily reflect the views or policies of the NIH, the Department of Health and Human Services, or the U.S. Government. Mention of trade names, commercial products, or organizations does not imply endorsement by the U.S. Government.

## Author Contributions

Conceptualization, D.T., Y.P.; methodology, D.T., F.E., Y.I., C.S., M.K.; investigation, D.T., Y.I., F.E., A.L., Y.P.; resources, Y.S.K., N.R.; formal analysis, D.T., Y.I., F.E., A.L., Y.P.; visualization, D.T., Y.I., F.E., Y.P.; supervision, A.L., M.I.A., Y.P.; writing– original draft, D.T., Y.I., Y.P.; and writing– review & editing, D.T., Y.I., A.L., M.I.A., and Y.P.

## Data Availability Statement

The single-cell RNA sequencing data generated in this study have been deposited in the Gene Expression Omnibus (GEO) under accession number GSE336003. Raw sequencing data are available through the Sequence Read Archive (SRA) under the associated accession. All other data supporting the findings of this study are available within the article and its Supplementary Information files.

## Supporting Information

**S1. *SLFN11*-correlated gene networks.**

Two-dimensional UMAP projections (as in Fig. 4) showing expression of gene lists in the indicated cell lines.

**S2. Distribution of SLFN11 expression in the remaining SCLC samples from the same single-cell cohort analyzed in Figure 6**.

Histograms showing the distribution of single-cell SLFN11 expression across the remaining 14 SCLC samples from the publicly available scRNA-seq dataset analyzed in Fig. 6. SLFN11 expression is shown as log_2_(X + 1), where X represents normalized transcript counts. Numbers in parentheses indicate the mean SLFN11 expression level for each sample. Samples are ordered according to decreasing mean SLFN11 expression.

## References

1. Mavrommatis E, Fish EN, Platanias LC. The schlafen family of proteins and their regulation by interferons. J Interferon Cytokine Res. 2013;33(4):206–10.

2. Schwarz DA, Katayama CD, Hedrick SM. Schlafen, a new family of growth regulatory genes that affect thymocyte development. Immunity. 1998;9(5):657–68.

3. Al-Marsoummi S, Vomhof-DeKrey EE, Basson MD. Schlafens: Emerging Proteins in Cancer Cell Biology. Cells. 2021;10(9).

4. Metzner FJ, Wenzl SJ, Kugler M, Krebs S, Hopfner KP, Lammens K. Mechanistic understanding of human SLFN11. Nat Commun. 2022;13(1):5464.

5. Bustos O, Naik S, Ayers G, Casola C, Perez-Lamigueiro MA, Chippindale PT, et al. Evolution of the Schlafen genes, a gene family associated with embryonic lethality, meiotic drive, immune processes and orthopoxvirus virulence. Gene. 2009;447(1):1–11.

6. Jo U, Murai Y, Agama KK, Sun Y, Saha LK, Yang X, et al. TOP1-DNA Trapping by Exatecan and Combination Therapy with ATR Inhibitor. Mol Cancer Ther. 2022;21(7):1090–102.

7. Murai J, Tang SW, Leo E, Baechler SA, Redon CE, Zhang H, et al. SLFN11 Blocks Stressed Replication Forks Independently of ATR. Mol Cell. 2018;69(3):371–84 e6.

8. Zoppoli G, Regairaz M, Leo E, Reinhold WC, Varma S, Ballestrero A, et al. Putative DNA/RNA helicase Schlafen-11 (SLFN11) sensitizes cancer cells to DNA-damaging agents. Proc Natl Acad Sci U S A. 2012;109(37):15030–5.

9. Murai J, Thomas A, Miettinen M, Pommier Y. Schlafen 11 (SLFN11), a restriction factor for replicative stress induced by DNA-targeting anti-cancer therapies. Pharmacol Ther. 2019;201:94–102.

10. Scattolin D, Maso AD, Ferro A, Frega S, Bonanno L, Guarneri V, et al. The emerging role of Schlafen-11 (SLFN11) in predicting response to anticancer treatments: Focus on small cell lung cancer. Cancer Treatment Reviews. 2024;128:102768.

11. Takashima T, Taniyama D, Sakamoto N, Yasumoto M, Asai R, Hattori T, et al. Schlafen 11 predicts response to platinum-based chemotherapy in gastric cancers. Br J Cancer. 2021;125(1):65–77.

12. Taniyama D, Sakamoto N, Takashima T, Takeda M, Pham QT, Ukai S, et al. Prognostic impact of Schlafen 11 in bladder cancer patients treated with platinum-based chemotherapy. Cancer Sci. 2022;113(2):784–95.

13. Akashi H, Yachida N, Ueda H, Yamaguchi M, Yamawaki K, Tamura R, et al. SLFN11 is a BRCA Independent Biomarker for the Response to Platinum-Based Chemotherapy in High-Grade Serous Ovarian Cancer and Clear Cell Ovarian Carcinoma. Mol Cancer Ther. 2024;23(1):106–16.

14. Chakraborty K, Burman R, Satheesh S, Kieffer M, Karuhatty C, Yuan ZF, et al. Targeting glycerophospholipid biosynthesis overcomes chemoresistance driven by SLFN11 loss in Ewing sarcoma. Cell Death Dis. 2026.

15. Croushore EE, Koppenhafer SL, Goss KL, Geary EL, Gordon DJ. Activator Protein-1 (AP-1) Signaling Inhibits the Growth of Ewing Sarcoma Cells in Response to DNA Replication Stress. Cancer Res Commun. 2023;3(8):1580–93.

16. Masuda K, Yoshida T, Motoi N, Shinno Y, Matsumoto Y, Okuma Y, et al. Schlafen 11 Expression in Patients With Small Cell Lung Cancer and Its Association With Clinical Outcomes. Thorac Cancer. 2025;16(1):e15529.

17. Nakata S, Murai J, Okada M, Takahashi H, Findlay TH, Malebranche K, et al. Epigenetic upregulation of Schlafen11 renders WNT- and SHH-activated medulloblastomas sensitive to cisplatin. Neuro Oncol. 2022.

18. Pietanza MC, Waqar SN, Krug LM, Dowlati A, Hann CL, Chiappori A, et al. Randomized, Double-Blind, Phase II Study of Temozolomide in Combination With Either Veliparib or Placebo in Patients With Relapsed-Sensitive or Refractory Small-Cell Lung Cancer. J Clin Oncol. 2018;36(23):2386–94.

19. Raynaud CM, Ahmed EI, Jabeen A, Sanchez A, Sherif S, Carneiro-Lobo TC, et al. Modulation of SLFN11 induces changes in DNA Damage response in breast cancer. Cancer Cell Int. 2023;23(1):291.

20. Wang HX, Zhao ZP, Du XY, Peng SL, Xu HY, Tang W, et al. SLFN11 promotes clear cell renal cell carcinoma progression via the PI3K/AKT signaling pathway. Med Oncol. 2024;41(2):54.

21. Willis SE, Winkler C, Roudier MP, Baird T, Marco-Casanova P, Jones EV, et al. Retrospective analysis of Schlafen11 (SLFN11) to predict the outcomes to therapies affecting the DNA damage response. Br J Cancer. 2021;125(12):1666–76.

22. Zhou C, Weng J, Liu C, Liu S, Hu Z, Xie X, et al. Disruption of SLFN11 deficiency-induced CCL2 signaling and macrophage M2 polarization potentiates anti-PD-1 therapy efficacy in hepatocellular carcinoma. Gastroenterology. 2023.

23. Kaczorowski M, Ylaya K, Chlopek M, Taniyama D, Pommier Y, Lasota J, et al. Immunohistochemical Evaluation of Schlafen 11 (SLFN11) Expression in Cancer in the Search of Biomarker-Informed Treatment Targets: A Study of 127 Entities Represented by 6658 Tumors. Am J Surg Pathol. 2024.

24. Takashima T, Sakamoto N, Murai J, Taniyama D, Honma R, Ukai S, et al. Immunohistochemical analysis of SLFN11 expression uncovers potential non-responders to DNA-damaging agents overlooked by tissue RNA-seq. Virchows Arch. 2021;478(3):569–79.

25. Tang SW, Bilke S, Cao L, Murai J, Sousa FG, Yamade M, et al. SLFN11 is a transcriptional target of EWS-FLI1 and a determinant of drug response in Ewing’s sarcoma. Clinical cancer research : an official journal of the American Association for Cancer Research. 2015;21(18):4184–93.

26. Murai Y, Jo U, Murai J, Fukuda S, Takebe N, Pommier Y. Schlafen 11 expression in human acute leukemia cells with gain-of-function mutations in the interferon-JAK signaling pathway. iScience. 2021;24(10):103173.

27. Borrego AR, Corona-Ayala C, Salvador JC, Valdez FC, Llano M. Gene Expression Regulation of the Type I Interferon-Induced Protein Schlafen 11. The FASEB Journal. 2020;34(S1):1–.

28. Nogales V, Reinhold WC, Varma S, Martinez-Cardus A, Moutinho C, Moran S, et al. Epigenetic inactivation of the putative DNA/RNA helicase SLFN11 in human cancer confers resistance to platinum drugs. Oncotarget. 2016;7(3):3084–97.

29. Tang SW, Bilke S, Cao L, Murai J, Sousa FG, Yamade M, et al. SLFN11 Is a Transcriptional Target of EWS-FLI1 and a Determinant of Drug Response in Ewing Sarcoma. Clin Cancer Res. 2015;21(18):4184–93.

30. Gardner EE, Lok BH, Schneeberger VE, Desmeules P, Miles LA, Arnold PK, et al. Chemosensitive Relapse in Small Cell Lung Cancer Proceeds through an EZH2-SLFN11 Axis. Cancer Cell. 2017;31(2):286–99.

31. Stewart CA, Gay CM, Xi Y, Sivajothi S, Sivakamasundari V, Fujimoto J, et al. Single-cell analyses reveal increased intratumoral heterogeneity after the onset of therapy resistance in small-cell lung cancer. Nat Cancer. 2020;1:423–36.

32. Kinker GS, Greenwald AC, Tal R, Orlova Z, Cuoco MS, McFarland JM, et al. Pan-cancer single-cell RNA-seq identifies recurring programs of cellular heterogeneity. Nat Genet. 2020;52(11):1208–18.

33. Puram SV, Tirosh I, Parikh AS, Patel AP, Yizhak K, Gillespie S, et al. Single-Cell Transcriptomic Analysis of Primary and Metastatic Tumor Ecosystems in Head and Neck Cancer. Cell. 2017;171(7):1611–24 e24.

34. Tirosh I, Izar B, Prakadan SM, Wadsworth MH, 2nd, Treacy D, Trombetta JJ, et al. Dissecting the multicellular ecosystem of metastatic melanoma by single-cell RNA-seq. Science. 2016;352(6282):189–96.

35. Patel AP, Tirosh I, Trombetta JJ, Shalek AK, Gillespie SM, Wakimoto H, et al. Single-cell RNA-seq highlights intratumoral heterogeneity in primary glioblastoma. Science. 2014;344(6190):1396–401.

36. Xiang X, Tao X, Hua K, Jiang H, Ding J. Single-cell RNA sequencing reveals tumor heterogeneity in small cell neuroendocrine cervical carcinoma. Commun Biol. 2025;8(1):184.

37. Hao Y, Hao S, Andersen-Nissen E, Mauck WM, 3rd, Zheng S, Butler A, et al. Integrated analysis of multimodal single-cell data. Cell. 2021;184(13):3573–87 e29.

38. van Dijk D, Sharma R, Nainys J, Yim K, Kathail P, Carr AJ, et al. Recovering Gene Interactions from Single-Cell Data Using Data Diffusion. Cell. 2018;174(3):716–29 e27.

39. Qiu P. Embracing the dropouts in single-cell RNA-seq analysis. Nat Commun. 2020;11(1):1169.

40. Subramanian A, Tamayo P, Mootha VK, Mukherjee S, Ebert BL, Gillette MA, et al. Gene set enrichment analysis: a knowledge-based approach for interpreting genome-wide expression profiles. Proc Natl Acad Sci U S A. 2005;102(43):15545–50.

41. Fang Z, Liu X, Peltz G. GSEApy: a comprehensive package for performing gene set enrichment analysis in Python. Bioinformatics. 2023;39(1).

42. Ragueneau E, Gong C, Sinquin P, Sevilla C, Beavers D, Grentner A, et al. The Reactome Knowledgebase 2026. Nucleic Acids Res. 2026;54(D1):D673–D81.

43. Liberzon A, Subramanian A, Pinchback R, Thorvaldsdottir H, Tamayo P, Mesirov JP. Molecular signatures database (MSigDB) 3.0. Bioinformatics. 2011;27(12):1739–40.

44. Chan JM, Quintanal-Villalonga A, Gao VR, Xie Y, Allaj V, Chaudhary O, et al. Signatures of plasticity, metastasis, and immunosuppression in an atlas of human small cell lung cancer. Cancer Cell. 2021;39(11):1479–96 e18.

45. Rajapakse VN, Luna A, Yamade M, Loman L, Varma S, Sunshine M, et al. CellMinerCDB for Integrative Cross-Database Genomics and Pharmacogenomics Analyses of Cancer Cell Lines. iScience. 2018;10:247–64.

46. Tlemsani C, Pongor L, Elloumi F, Girard L, Huffman KE, Roper N, et al. SCLC-CellMiner: A Resource for Small Cell Lung Cancer Cell Line Genomics and Pharmacology Based on Genomic Signatures. Cell Rep. 2020;33(3):108296.

47. Elloumi F, Reinhold WC, Varma S, Wang Y, Kinali M, Arakawa Y, et al. CellMiner cross-database (CellMinerCDB) version 2.2 for explorations of patient-derived cancer cell line pharmacogenomics. Nucleic Acids Res. 2025.

48. Barretina J, Caponigro G, Stransky N, Venkatesan K, Margolin AA, Kim S, et al. The Cancer Cell Line Encyclopedia enables predictive modelling of anticancer drug sensitivity. Nature. 2012;483(7391):603–307.

49. Garnett MJ, Edelman EJ, Heidorn SJ, Greenman CD, Dastur A, Lau KW, et al. Systematic identification of genomic markers of drug sensitivity in cancer cells. Nature. 2012;483(7391):570–5.

50. Khandagale P, Welter AL, Huang SN, Taniyama D, Machida Y, Pommier Y. SLFN11 puts the brakes on Alternative lengthening of telomeres. bioRxiv. 2025.

51. Murai J, Feng Y, Yu GK, Ru Y, Tang SW, Shen Y, et al. Resistance to PARP inhibitors by SLFN11 inactivation can be overcome by ATR inhibition. Oncotarget. 2016;7(47):76534–50.

52. Lahav G, Rosenfeld N, Sigal A, Geva-Zatorsky N, Levine AJ, Elowitz MB, et al. Dynamics of the p53-Mdm2 feedback loop in individual cells. Nat Genet. 2004;36(2):147–50.

53. Batchelor E, Mock CS, Bhan I, Loewer A, Lahav G. Recurrent initiation: a mechanism for triggering p53 pulses in response to DNA damage. Mol Cell. 2008;30(3):277–89.

54. Sun F, Das MS. Targeting the EZH2-SLFN11 pathway-a lesson in developing molecularly-informed treatments for recurrent small cell lung cancer. Transl Cancer Res. 2024;13(12):6608–12.

55. Vanesa Nogales WCR, Sudhir Varma, Anna Martinez-Cardus, Catia Moutinho, Sebastian Moran, Holger Heyn, Ana Sebio, Agusti Barnadas, Yves Pommier, Manel Esteller. Epigenetic inactivation of the putative DNA:RNA helicase SLFN11 in human cancer confers resistance to platinum drugs.pdf. 2016.

56. Tang SW, Thomas A, Murai J, Trepel JB, Bates SE, Rajapakse VN, et al. Overcoming Resistance to DNA-Targeted Agents by Epigenetic Activation of Schlafen 11 (SLFN11) Expression with Class I Histone Deacetylase Inhibitors. Clin Cancer Res. 2018;24(8):1944–53.

57. Murai Y, Jo U, Murai J, Jenkins LM, Huang SN, Chakka S, et al. SLFN11 Inactivation Induces Proteotoxic Stress and Sensitizes Cancer Cells to Ubiquitin Activating Enzyme Inhibitor TAK-243. Cancer Res. 2021;81(11):3067–78.

58. Barretina J, Caponigro G, Stransky N, Venkatesan K, Margolin AA, Kim S, et al. The Cancer Cell Line Encyclopedia enables predictive modelling of anticancer drug sensitivity. Nature. 2012;483(7391):603–7.

59. Murai Y, Jo U, Murai J, Jenkins L, Huang SN, Chakka S, et al. SLFN11 inactivation induces proteotoxic stress and sensitizes cancer cells to ubiquitin activating enzyme inhibitor TAK-243. Cancer Res. 2021;81(11):3067–78.

60. Ogawa A, Izumikawa K, Tate S, Isoyama S, Mori M, Fujiwara K, et al. SLFN11-mediated ribosome biogenesis impairment induces TP53-independent apoptosis. Mol Cell. 2025;85(5):894–912 e10.

61. Scattolin D, Heijboer FWJ, Derks JL, Thüsen JHV, Dumoulin DW, Speel EJM, et al. Clinical insight on the pathway of SLFN11 as emergent biomarker in SCLC. Critical Reviews in Oncology Hematology. 2025;215.

